# Identification of gingipains in glioblastoma tumors and evidence that *P. gingivalis* infection drives IL-6 and PD-L1 expression in glioma cells

**DOI:** 10.1101/2025.11.13.686868

**Authors:** Ella M. Moore, Laurent A. Bekale, Zin Mie Mie Tun, Jing Chen, Mark I. Ryder, Florian Ermini, Stephen S. Dominy, Annelise E. Barron

**Affiliations:** Department of Bioengineering, Schools of Engineering & of Medicine, Stanford University, Stanford, California, 94305, USA

## Abstract

Glioblastoma multiforme (GBM), a highly aggressive brain tumor that accounts for approximately 60% of all gliomas and 48% of primary central nervous system malignancies, is incurable and poorly understood, with a median survival of only 15 months after diagnosis. Thus, there is an urgent need to understand GBM pathogenesis in order to develop an effective treatment. Recent research has revealed frequent Alzheimer’s disease (AD) pathology in the brains of patients with GBM, *i.e.*, amyloid beta (Aβ) and hyperphosphorylated tau (pTau), indicating that GBM and AD may share some unknown environmental risk. Since chronic periodontitis (CP), and specifically *Porphyromonas gingivalis* (*P. gingivalis)*, a keystone bacterial pathogen in CP, have emerged as risk factors for both AD and GBM, we investigated whether *P. gingivalis* gingipain virulence factors could be identified in GBM tissue samples and whether *P. gingivalis* infection affects glioma cell behavior. Using immunohistochemistry on tissue microarrays (70 GBM cores from 35 patients; 34 cerebral tissue cores from 17 patients), we quantified the presence of arginine-gingipain B (RgpB) and lysine-gingipain (Kgp) antigens. Both gingipains showed significantly elevated staining in GBM samples compared to controls (**p < .01, ****p < .0001, respectively), with Kgp levels notably higher than RgpB within GBM tissue (****p < .0001). In functional assays using U251 glioma cells, *P. gingivalis* infection induced robust, dose-dependent IL-6 secretion (peaking at MOI 5), increased PD-L1 expression by 30% (*p = .036), and significantly enhanced cell invasiveness (**p < .01) in a viability-dependent manner. These findings demonstrate that *P. gingivalis* gingipains are present at elevated levels in GBM tissue and that *P. gingivalis* infection reprograms glioma cells to adopt an immunosuppressive, invasive phenotype through upregulation of the IL-6/PD-L1 axis, suggesting a potential microbial contribution to GBM pathogenesis and immune evasion.

**Key points:**

- Glioblastoma multiforme (GBM) patients have frequent Alzheimer’s disease (AD) neuropathological changes in the tumor-adjacent cortex, indicating that GBM tumors may share some environmental risk factors with AD.
- This study identifies gingipain antigens in GBM tissue samples at significantly elevated levels compared to healthy controls, suggesting that *P. gingivalis* infection may be an environmental risk factor for both AD and GBM.
- In *in vitro* experiments, *P. gingivalis* infection of the human glioma cell line U251 upregulated IL-6 secretion and PD-L1 expression, and significantly increased cell invasiveness compared to uninfected cells.

## Introduction

In 2010, based on findings of a significant positive correlation between Alzheimer’s disease (AD) and malignant brain tumor mortality rate, a hypothesis was published that stated glioblastoma and dementia may share a common cause through “an as yet unknown peripheral tissue pathway that can promote the progression of both diseases^1^.” The author suggested that identification of the common pathway could lead to new treatments for both glioblastoma multiforme (GBM) and AD.

Since then, a number of studies have demonstrated the presence of amyloid beta (Aβ) and phosphorylated tau (pTau), pathologic hallmarks of AD, in and around GBM tumors^2,3^. Aβ has been shown to be an antimicrobial peptide in the innate immune response against invading pathogens^4^, and its association with GBM suggests the possibility of an infectious agent(s) in or around the tumor. For example, cytomegalovirus (CMV) has been identified in human malignant gliomas and has been a focus of research in possible viral causes of GBM^5,6^.

Recently, a bacterial pathogen, *Porphyromonas gingivalis* (*P. gingivalis*), best known for its role as a keystone pathogen in chronic periodontitis (CP)^7^, has emerged as a possible infectious agent in GBM pathogenesis^8^. *P. gingivalis* is a gram-negative, anaerobic, and asaccharolytic bacterium capable of invading cells and persisting intracellularly^9^. In a first-of-a-kind study, patients with glioma underwent periodontal examinations prior to glioma surgery, and their tumor biopsy samples were measured for Ki-67, a tumor proliferation marker^8^. Patients with evidence of more advanced periodontal disease had a significantly higher tumor Ki-67 labeling index, indicating a larger proportion of glioma tumor cells were dividing. *In vitro, P. gingivalis* LPS promoted glioma cell proliferation and migration by activating the pro-tumor Akt pathway, supporting an association between *P. gingivalis* periodontal infection and glioma progression^8^.

*P. gingivalis* has also been linked to the pathogenesis of AD. Chronic oral administration of *P. gingivalis* in wild-type mice showed that *P. gingivalis* and/or its gingipain virulence factors could translocate to the brain and produce neuropathology that is consistent with AD, including the production of Aβ42 and pTau^10^. Using IHC, gingipains were shown to localize intranuclearly and perinuclearly in microglia, astrocytes, and neurons. Neurodegeneration was evident in the *P. gingivalis*-infected group but not in sham-infected control mice^10^. Members of our team detected *P. gingivalis* gingipains in the middle temporal gyrus and hippocampus of human AD brains by immunohistochemistry, and their levels were correlated with AD diagnosis and with tau and ubiquitin pathology^11^. We also demonstrated in wild-type mice that oral *P. gingivalis* infection led to *P. gingivalis* brain invasion and Aβ42 induction, effects that could be blocked by small-molecule gingipain inhibitors. In an *in vitro* study, Aβ42 was shown to localize to the surface of *P. gingivalis* and kill the pathogen, reinforcing the antimicrobial function of Aβ42^11^.

The gingipains of *P. gingivalis* are cysteine protease virulence factors that are transported to the surface of the bacterium and can be packaged in outer membrane vesicles (OMV) that are released into the bloodstream and carried throughout the body^12^. Lysine-gingipain (Kgp) cleaves proteins on the C-terminal side of lysine residues, and arginine-gingipain B (RgpB) and arginine-gingipain A (RgpA) cleave proteins on the C-terminal side of arginine residues. Gingipains are essential for *P. gingivalis* survival and pathogenicity by inactivating host defenses, accomplishing iron and nutrient acquisition, and facilitating tissue destruction^9,13^.

Gingipains have also been connected to *P. gingivalis’s* oncogenic properties. Experiments have shown that *P. gingivalis* infection of macrophages upregulates the expression of Programmed death-ligand 1 (PD-L1) and that gingipains regulate PD-L1 isoform switches that favor enhanced PD-L1 docking with Programmed cell death protein 1 (PD-1), an immune checkpoint molecule expressed on T cells^14^. PD-L1 has been extensively studied and targeted in the cancer field due to its ability to suppress immune responses^15^. A recent study demonstrated that *in vitro, P. gingivalis* infection of dendritic cells “robustly” enhanced PD-L1 expression in a gingipain-dependent manner^15^. In a follow-on study using a syngeneic mouse oral cancer model, *P. gingivalis* infection significantly enhanced PD-L1 expression on dendritic cells from intratumoral tissues and exacerbated oral cancer progression, but this effect was not seen when mice were infected with a *P. gingivalis* lysine-gingipain (Kgp)-defective mutant^15^. In addition, *P. gingivalis* infection has been shown to upregulate PD-L1 in human colon cancer cells^16^ and prostate cancer cells^17^. In GBM tumors, a high abundance of tumor-associated macrophages (TAMs) has been identified with upregulated PD-L1 that suppresses CD8^+^ T cell function, limiting the clinical efficacy of PD-1/PD-L1 blockade^18^. It is currently unknown if a proportion of TAMs in GBM tumors are *P. gingivalis-*infected and/or contain gingipains that could drive PD-L1 overexpression.

A second important *P. gingivalis* oncogenic mechanism is non-canonical activation of beta-catenin (β-catenin) by gingipain proteolytic activity^19^. β-catenin signaling is a major pathway in the control of cell proliferation and tumorigenesis, and aberrant activation has been linked to carcinogenesis and tumor progression of several cancers^20^. β-catenin has been found to be commonly overexpressed in GBM^21^.

We hypothesized that *P. gingivalis* infection contributes to GBM pathogenesis by secreting gingipains to promote immunosuppression and cancer cell proliferation. We found that gingipain immunoreactivity was significantly elevated in GBM tissue samples compared to controls, with Kgp staining levels notably higher than RgpB. *In vitro* studies of *P. gingivalis* infection of human astrocytes and a glioma cell line demonstrated increased expression of interleukin-6 (IL-6), a cytokine linked to immunosuppression in the GBM microenvironment^22^. We also show that *P. gingivalis* infection of glioma cells increased PD-L1 expression and invasiveness compared to uninfected cells. Our results indicate that *P. gingivalis* infection may be a peripheral tissue pathway^1^ that can promote the progression of both glioma and AD.

## Results

### Gingipain antigens are increased in glioblastoma tissue

Emerging evidence indicates that GBM and dementia may share similar disease mechanisms. To investigate whether *P. gingivalis* could be a common microbial factor contributing to both conditions, 34 cerebral tissue cores from 17 neurologically normal individuals and 70 tumor tissue cores from 35 GBM patients were analyzed using immunohistochemistry with antibodies specific to the *P. gingivalis* gingipains Kgp and RgpB. The demographic and clinical details of the GBM and control donors are summarized in **Tables 1 and 2**, respectively.

**Table 1:** Demographic, clinical characteristics, and gingipain immunostaining results of the GBM tissue samples. Patient ID was assigned according to the placement of the core in the microarray. Average age is 44.5 years. Samples are 54.3% female, 45.7% male. Net CAB101 depicts the net area (%) stained for RgpB, and Net CAB102 depicts the net area (%) stained for Kgp. Net area stained is the percentage stained for the respective primary antibody subtracted by the percentage stained on the control sample.

| Microarray | Patient ID | Sample | Age | Sex | Net CAB 101 | Net CAB 102 | Pathology Diagnosis |
| --- | --- | --- | --- | --- | --- | --- | --- |
| GL805 | 1 | A1 | 59 | F | -0.17% | 0.38% | Glioblastoma |
| GL805 | 1 | A2 | 59 | F | -0.15% | 12.48% | Glioblastoma |
| GL805 | 2 | A3 | 59 | M | 0.12% | 44.80% | Glioblastoma |
| GL805 | 2 | A4 | 59 | M | 1.01% | 37.39% | Glioblastoma |
| GL805 | 3 | A5 | 33 | F | 0.34% | 80.38% | Glioblastoma |
| GL805 | 3 | A6 | 33 | F | 1.13% | 94.02% | Glioblastoma |
| GL805 | 4 | A7 | 47 | M | 1.01% | 39.01% | Glioblastoma |
| GL805 | 4 | A8 | 47 | M | 1.71% | 19.55% | Glioblastoma |
| GL805 | 5 | A9 | 9 | M | 0.86% | 75.23% | Glioblastoma |
| GL805 | 5 | A10 | 9 | M | 3.41% | 66.01% | Glioblastoma |
| GL805 | 6 | B1 | 64 | F | 8.13% | 16.54% | Glioblastoma |
| GL805 | 6 | B2 | 64 | F | 1.22% | 15.97% | Glioblastoma |
| GL805 | 7 | B3 | 17 | M | 7.74% | 78.44% | Glioblastoma |
| GL805 | 7 | B4 | 17 | M | 10.64% | 72.34% | Glioblastoma |
| GL805 | 8 | B5 | 21 | M | 7.62% | 84.21% | Glioblastoma |
| GL805 | 8 | B6 | 21 | M | 7.34% | 72.19% | Glioblastoma |
| GL805 | 9 | B7 | 70 | F | 2.91% | 96.55% | Glioblastoma |
| GL805 | 9 | B8 | 70 | F | 2.81% | 96.90% | Glioblastoma |
| GL805 | 10 | B9 | 46 | M | 14.47% | 78.24% | Glioblastoma |
| GL805 | 10 | B10 | 46 | M | 6.35% | 83.62% | Glioblastoma |
| GL805 | 11 | C1 | 32 | F | 8.33% | 42.88% | Glioblastoma |
| GL805 | 11 | C2 | 32 | F | 7.68% | 92.38% | Glioblastoma |
| GL805 | 12 | C3 | 32 | M | 1.31% | 71.57% | Glioblastoma |
| GL805 | 12 | C4 | 32 | M | 1.53% | 81.89% | Glioblastoma |
| GL805 | 13 | C5 | 48 | F | 3.82% | 49.41% | Glioblastoma |
| GL805 | 13 | C6 | 48 | F | 1.40% | 80.64% | Glioblastoma |
| GL805 | 14 | C7 | 50 | M | 2.34% | 77.32% | Glioblastoma |
| GL805 | 14 | C8 | 50 | M | 1.47% | 75.85% | Glioblastoma |
| GL805 | 15 | C9 | 59 | M | 4.15% | 69.31% | Glioblastoma |
| GL805 | 15 | C10 | 59 | M | 0.51% | 81.74% | Glioblastoma |
| GL805 | 16 | D1 | 59 | F | 1.64% | 13.96% | Glioblastoma |
| GL805 | 16 | D2 | 59 | F | 4.43% | 49.33% | Glioblastoma |
| GL805 | 17 | D3 | 16 | F | 2.05% | 70.28% | Glioblastoma |
| GL805 | 17 | D4 | 16 | F | 1.95% | 69.07% | Glioblastoma |
| GL805 | 18 | D5 | 40 | F | 2.93% | 61.07% | Glioblastoma |
| GL805 | 18 | D6 | 40 | F | 2.71% | -0.22% | Glioblastoma |
| GL805 | 19 | D7 | 80 | M | 12.83% | 89.86% | Glioblastoma |
| GL805 | 19 | D8 | 80 | M | 5.84% | 84.66% | Glioblastoma |
| GL805 | 20 | D9 | 31 | M | 0.23% | 84.85% | Glioblastoma |
| GL805 | 20 | D10 | 31 | M | 0.58% | 84.67% | Glioblastoma |
| GL805 | 21 | E1 | 45 | M | 5.79% | 47.12% | Glioblastoma |
| GL805 | 21 | E2 | 45 | M | 12.04% | 40.92% | Glioblastoma |
| GL805 | 22 | E3 | 46 | F | 4.48% | 63.56% | Glioblastoma |
| GL805 | 22 | E4 | 46 | F | 16.17% | 53.53% | Glioblastoma |
| GL805 | 23 | E5 | 41 | F | 17.30% | 79.71% | Glioblastoma |
| GL805 | 23 | E6 | 41 | F | 2.32% | 55.08% | Glioblastoma |
| GL805 | 24 | E7 | 45 | M | 3.65% | 44.53% | Glioblastoma |
| GL805 | 24 | E8 | 45 | M | 3.10% | 47.72% | Glioblastoma |
| GL805 | 25 | E9 | 43 | M | 3.08% | 83.87% | Glioblastoma |
| GL805 | 25 | E10 | 43 | M | 10.68% | 80.15% | Glioblastoma |
| GL805 | 26 | F1 | 64 | F | 44.04% | 36.67% | Glioblastoma |
| GL805 | 26 | F2 | 64 | F | 3.41% | 78.96% | Glioblastoma |
| GL805 | 27 | F3 | 52 | F | 0.35% | 62.01% | Glioblastoma |
| GL805 | 27 | F4 | 52 | F | 0.97% | 79.37% | Glioblastoma |
| GL805 | 28 | F5 | 57 | M | 0.22% | 53.35% | Glioblastoma |
| GL805 | 28 | F6 | 57 | M | 0.14% | 65.27% | Glioblastoma |
| GL805 | 29 | F7 | 8 | F | 71.64% | 44.93% | Glioblastoma |
| GL805 | 29 | F8 | 8 | F | 78.08% | 53.35% | Glioblastoma |
| GL805 | 30 | F9 | 67 | F | 4.94% | 71.54% | Glioblastoma |
| GL805 | 30 | F10 | 67 | F | 22.94% | 51.62% | Glioblastoma |
| GL805 | 31 | G1 | 22 | F | 0.25% | 5.38% | Glioblastoma |
| GL805 | 31 | G2 | 22 | F | 2.69% | 84.23% | Glioblastoma |
| GL805 | 32 | G3 | 47 | F | 44.58% | 45.34% | Glioblastoma |
| GL805 | 32 | G4 | 47 | F | 58.42% | 21.64% | Glioblastoma |
| GL805 | 33 | G5 | 56 | M | 0.71% | 33.96% | Glioblastoma |
| GL805 | 33 | G6 | 56 | M | 1.21% | 48.77% | Glioblastoma |
| GL805 | 34 | G7 | 44 | F | 0.84% | 13.43% | Glioblastoma |
| GL805 | 34 | G8 | 44 | F | 0.80% | 25.74% | Glioblastoma |
| GL805 | 35 | G9 | 51 | F | 1.58% | 83.90% | Glioblastoma |
| GL805 | 35 | G10 | 51 | F | 2.50% | 78.95% | Glioblastoma |

**Table 2:** Demographic and clinical characteristics, and gingipain immunostaining results of the control brain tissue samples. Patient ID was assigned according to the order of the cores in the microarray. Average age is 39.9 years. Samples are 23.5% female, 76.47% male. Net CAB101 depicts the net area (%) stained for RgpB, and Net CAB102 depicts the net area (%) stained for Kgp. Net area stained is the percentage stained for the respective primary antibody subtracted by the percentage stained on the control sample.

| Microarray | Patient ID | Sample | Age | Sex | Net CAB 101 | Net CAB 102 | Pathology Diagnosis |
| --- | --- | --- | --- | --- | --- | --- | --- |
| GLN801 | 1 | A1 | 54 | F | 0.04% | 0.12% | Cerebrum gray matter |
| GLN801 | 1 | A2 | 54 | F | 0.04% | 0.18% | Cerebrum gray matter |
| GLN801 | 2 | A3 | 45 | M | 0.08% | 20.83% | Cerebrum white matter |
| GLN801 | 2 | A4 | 45 | M | 0.07% | 6.85% | Cerebrum white matter |
| GLN801 | 3 | A5 | 23 | M | 0.17% | 40.69% | Cerebrum gray matter |
| GLN801 | 3 | A6 | 23 | M | 0.14% | 41.58% | Cerebrum gray matter |
| GLN801 | 4 | A7 | 54 | F | 0.04% | 0.35% | Cerebrum gray matter |
| GLN801 | 4 | A8 | 54 | F | -0.08% | 0.03% | Cerebrum gray matter |
| GLN801 | 5 | A9 | 45 | M | 0.45% | 6.95% | Cerebrum gray matter |
| GLN801 | 5 | A10 | 45 | M | -0.09% | 17.33% | Cerebrum gray matter |
| GLN801 | 6 | A11 | 23 | M | -0.02% | 2.43% | Cerebrum gray matter |
| GLN801 | 6 | A12 | 23 | M | 0.02% | 4.90% | Cerebrum gray matter |
| GLN801 | 7 | B1 | 31 | M | 0.13% | 5.59% | Cerebrum gray matter |
| GLN801 | 7 | B2 | 31 | M | 0.36% | 78.56% | Cerebrum gray matter |
| GLN801 | 8 | B3 | 45 | M | 0.05% | 14.92% | Cerebrum white matter |
| GLN801 | 8 | B4 | 45 | M | 0.09% | 7.21% | Cerebrum white matter |
| GLN801 | 9 | B5 | 23 | M | 0.59% | 42.72% | Cerebrum gray matter |
| GLN801 | 9 | B6 | 23 | M | 2.77% | 63.54% | Cerebrum gray matter |
| GLN801 | 10 | B7 | 45 | M | 0.94% | 72.74% | Cerebrum gray matter |
| GLN801 | 10 | B8 | 45 | M | 0.32% | 57.81% | Cerebrum gray matter |
| GLN801 | 11 | B9 | 23 | M | 0.07% | 20.88% | Cerebrum gray matter |
| GLN801 | 11 | B10 | 23 | M | 0.14% | 36.57% | Cerebrum gray matter |
| GLN801 | 12 | B11 | 27 | M | 0.09% | 11.58% | Cerebrum gray matter |
| GLN801 | 12 | B12 | 27 | M | 0.44% | 9.38% | Cerebrum gray matter |
| GL805 | 36 | H1 | 58 | F | 0.61% | 25.95% | Cerebral tissue |
| GL805 | 36 | H2 | 58 | F | 1.36% | 46.37% | Cerebral tissue |
| GL805 | 37 | H3 | 38 | M | 0.20% | 60.10% | Cerebral tissue |
| GL805 | 37 | H4 | 38 | M | 0.22% | 73.74% | Cerebral tissue |
| GL805 | 38 | H5 | 72 | M | 0.93% | 43.82% | Cerebral tissue |
| GL805 | 38 | H6 | 72 | M | 0.34% | 47.16% | Cerebral tissue |
| GL805 | 39 | H7 | 34 | M | 1.41% | 18.88% | Cerebral tissue |
| GL805 | 39 | H8 | 34 | M | 1.60% | 13.35% | Cerebral tissue |
| GL805 | 40 | H9 | 38 | F | 0.54% | 28.07% | Cerebral tissue |
| GL805 | 40 | H10 | 38 | F | 0.45% | 33.64% | Cerebral tissue |

Staining of both Kgp and RgpB antigens was significantly more prevalent in GBM tumor tissues compared to cerebral tissues from neurologically normal individuals (**Figure 1A**). Quantitative image analysis confirmed a significant increase in the gingipain-positive area (%) per core in GBM samples (**p < .01, ****p < .0001, respectively; unpaired two-tailed t-tests) (**Figures 1B–C**). Within GBM cores, Kgp staining intensity and prevalence were notably greater than those of RgpB (****p < .0001, unpaired two-tailed t-test) (**Figure 1D**). The relative distribution of Kgp and RgpB staining in GBM samples differed from patterns previously reported in AD brains, where both gingipains showed similarly high levels of immunoreactivity^11^. This distinct gingipain signature suggests that while both diseases may involve *P. gingivalis*-associated pathology, differential Kgp and RgpB levels could reflect disease-specific host–pathogen interactions and divergent down-stream pathogenic pathways.

**Figure 1:**
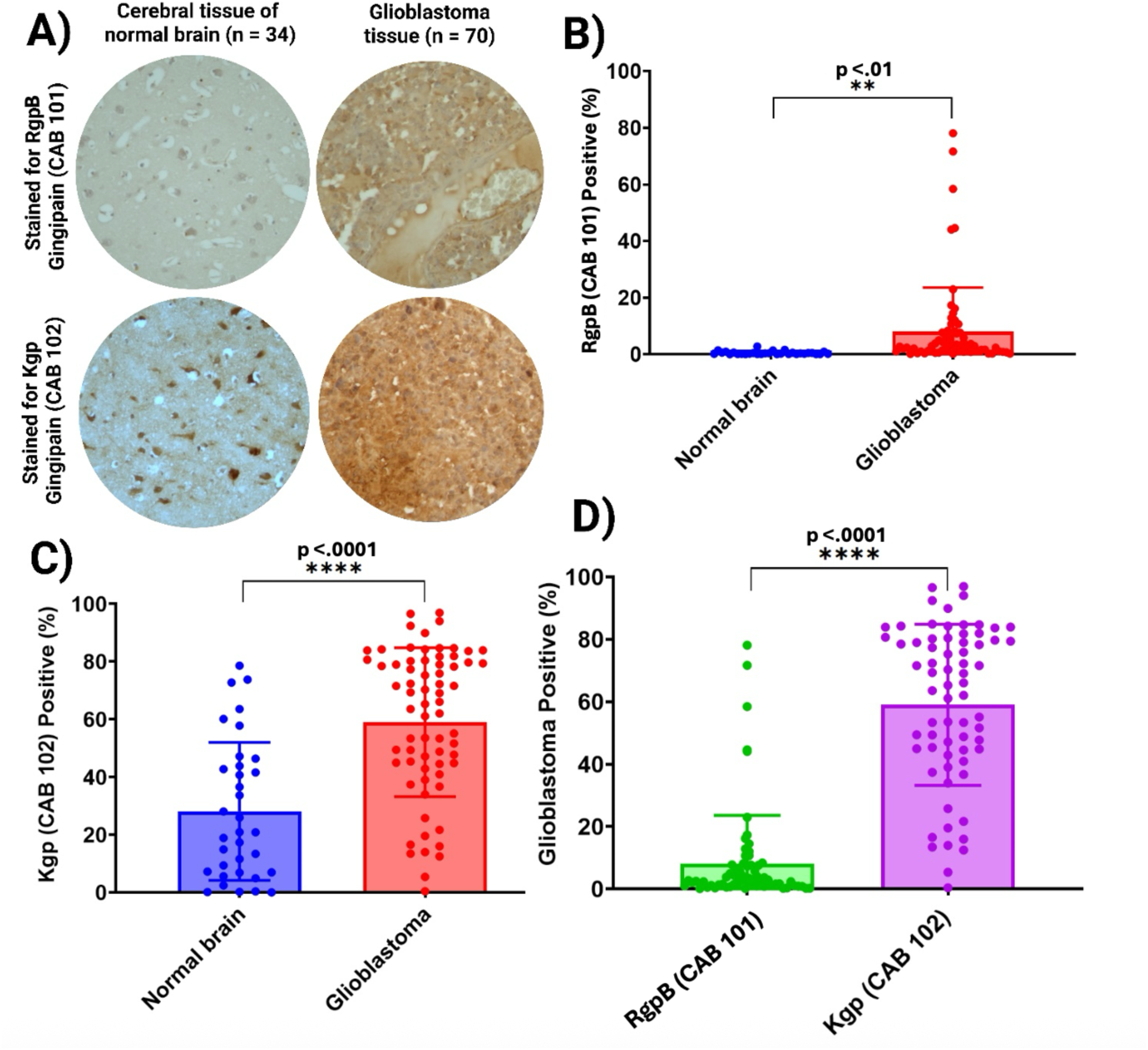
Gingipain antigens are enriched in glioblastoma tissue. **(A)** Representative immunohistochemistry (IHC) for RgpB (CAB101; top) and Kgp (CAB102; bottom) in cerebral tissue of normal brain (left; n = 34 cores) and GBM (right; n = 70 cores). Brown DAB indicates positive staining; hematoxylin counterstain. **(B)** Quantification of RgpB-positive area (%) per core in normal brain versus glioblastoma shows a significant increase in glioblastoma (**p < .01; unpaired two-tailed t-test). **(C)** Quantification of Kgp-positive area (%) per core in normal brain versus glioblastoma also shows a significant increase in glioblastoma (****p < .0001; unpaired two-tailed t-test). **(D)** Within glioblastoma cores, Kgp presence is higher than RgpB presence (****p < .0001; unpaired two-tailed t-test). Bars indicate mean ± SD; each dot represents one tissue core. *CAB101, anti-RgpB; CAB102, anti-Kgp. n values reflect independent tissue cores on the microarrays*.

Magnified images of stained tissue and control tissue are shown in **Figure 2**. In the first row, samples from a 9-year-old patient with GBM exhibit high Kgp antigen presence and minimal RgpB presence. In the second row, samples from an 8-year-old patient with GBM exhibit high antigen presence of both Kgp and RgpB. Rows three and four exhibit adult GBM patients (40-year-old and 80-year-old, respectively) with significant antigen presence of Kgp and minimal RgpB antigen presence, though slight presence of RgpB antigen can be seen in the 80-year-old subject. The last row exhibits samples from a neurologically normal control patient with minimal antigen presence of Kgp and RgpB (**Figure 2**).

**Figure 2:**
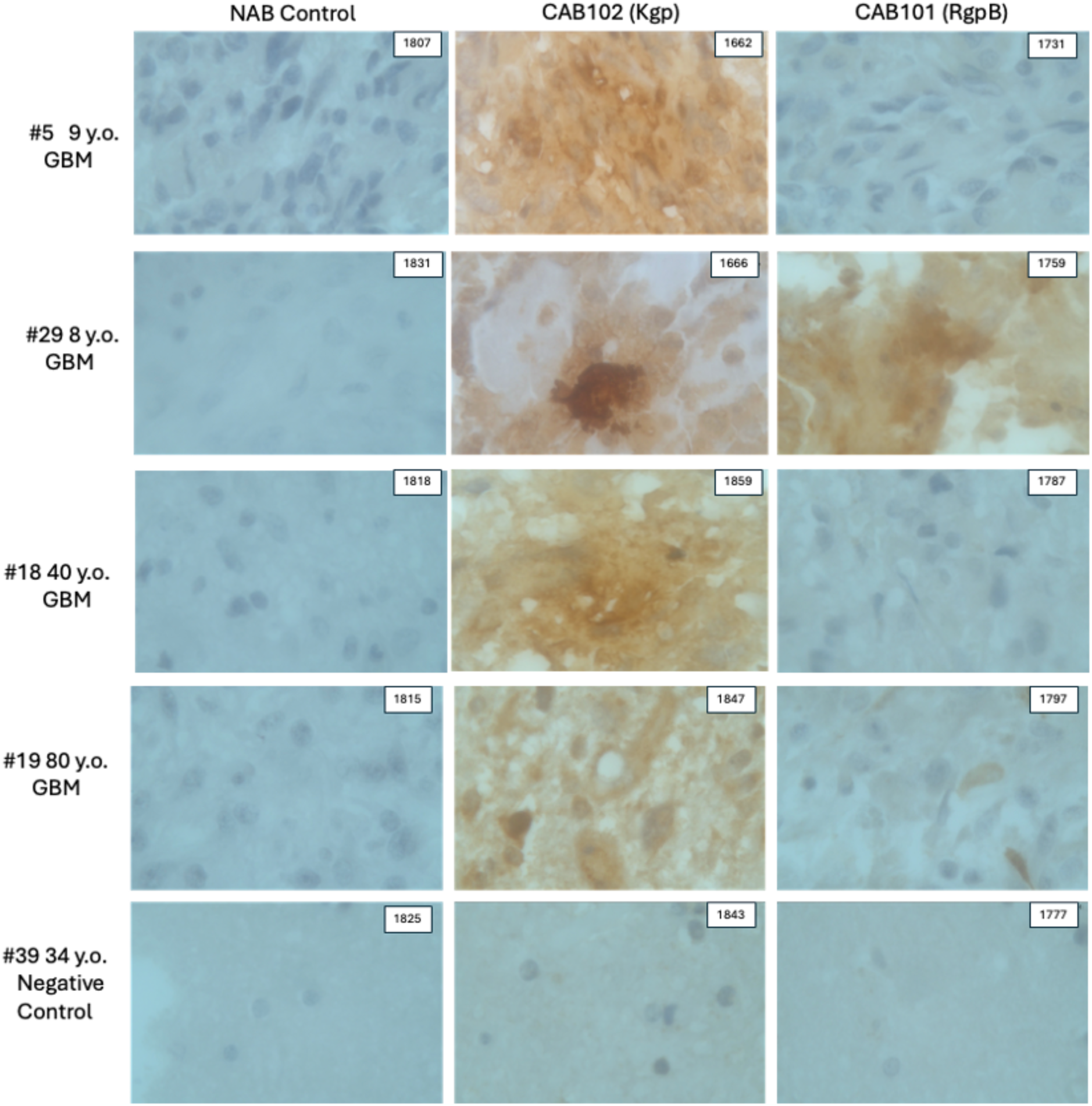
Glioblastoma brain tumor samples from pediatric and adult cases stained for *P. gingivalis* gingipains with CAB102 (anti-Kgp) and CAB101 (anti-RgpB). Formalin-fixed par-affin-embedded tissue sections were stained by immunohistochemistry using DAB chromogen (brown) with hematoxylin counterstain (blue). Four GBM cases (#5, #29, #18, #19) show positive staining with CAB102 and variable staining with CAB101, while the negative control (#39) shows no reactivity. NAB Control indicates that no primary antibody was applied. Numbers in upper right-hand corner of images correspond to the name of each image file. Images were taken at 100x magnification.

### P. gingivalis induces IL-6 production and upregulates PD-L1 expression in U251 glioma cells

Previous studies have shown that GBM–derived interleukin-6 (IL-6) acts as a key immunoregulatory cytokine that promotes tumor progression by driving the differentiation of immunosuppressive myeloid cells and increasing PD-L1 expression on tumor and stromal cells, thereby enabling immune evasion^23^. Given the established link between *P. gingivalis* infection and chronic inflammatory diseases, we hypothesized that *P. gingivalis* might activate a similar IL-6/PD-L1 signaling pathway in glioma cells. To test this, we measured IL-6 secretion and cell viability in U251 glioma cells and immortalized human astrocytes (HA) after a single 24 h exposure with increasing multiplicities of infection (MOI; baseline = 10^6^ CFU/mL) of *P. gingivalis*. Viability tests showed that astrocyte survival decreased at MOIs of 10 or higher, while U251 cells tolerated infection much better, with minimal viability loss even at high bacterial loads (**Figures 3A-B**). Quantifying IL-6 in the culture supernatants revealed a strong, dose-dependent cytokine response in both cell types, with IL-6 secretion peaking at MOI 2.5 in HA and MOI 5 in U251 cells (**Figures 3C-D**). At higher MOIs, IL-6 production declined, coinciding with cytotoxicity in HA cells and suggesting feedback inhibition or immune desensitization mechanisms in U251 cells. These results indicate that *P. gingivalis* can directly stimulate IL-6 production in both astrocytic and glioma cells, consistent with its role as a potent inflammatory stimulus capable of reprogramming host cell cytokine networks.

**Figure 3:**
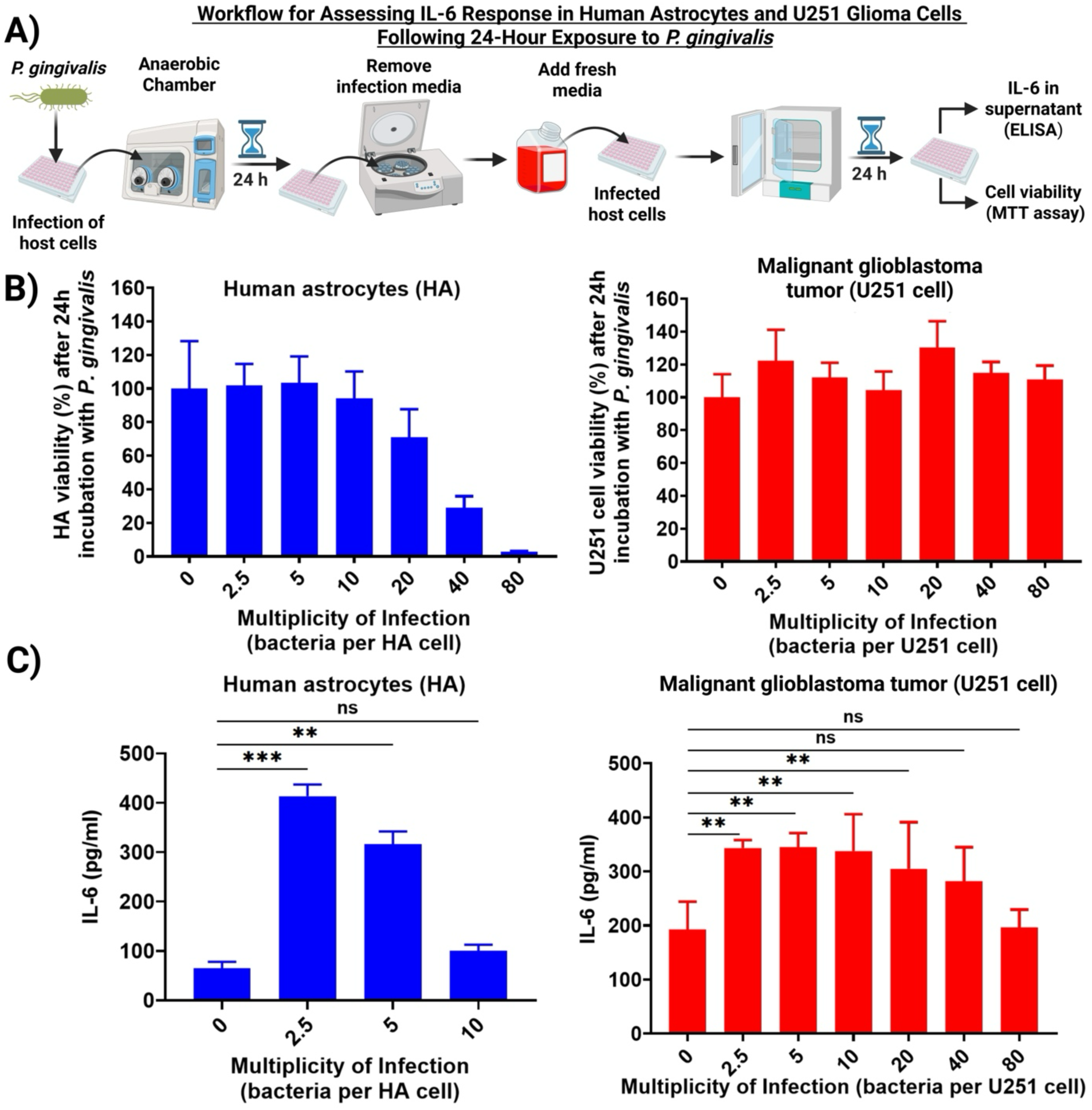
*P. gingivalis* selectively impairs astrocyte viability and induces IL-6 production in a dose-dependent manner. **(A)** Schematic of the experimental workflow used to assess IL-6 se-cretion and cell viability in human astrocytes (HA) and U251 glioma cells following a single 24-hour exposure to *P. gingivalis*. **(B)** Cell viability of HA (left) and U251 glioma cells (right) after 24 h infection with increasing multiplicities of infection (MOIs) of *P. gingivalis*. Astrocyte viabil-ity declined at MOIs ≥10, while U251 cells remained largely unaffected. **(C)** After infection, IL-6 levels in culture supernatants of HA (left) and U251 cells (right). Both cell types exhibited robust IL-6 secretion at low to moderate MOIs, with peak responses at MOI 2.5 (HA) and MOI 5 (U251). IL-6 production declined at higher MOIs, correlating with cytotoxicity in HA and possible immune suppression or feedback inhibition in U251 cells. Data represent mean ± SD. ns = not significant; **p < .01, ***p < .001 (unpaired two-tailed t-test).

Because IL-6 signaling is known to induce PD-L1 expression in GBM, we next examined whether *P. gingivalis* infection could directly upregulate PD-L1 protein levels in glioma cells. U251 cells were exposed to vehicle or *P. gingivalis* for 24 h, and whole-cell lysates were analyzed by immunoblot using an anti–PD-L1 antibody (NBP1-76769). Multiple bands were detected, indicating the presence of different PD-L1 glycoforms and isoforms, while GAPDH served as a loading control (**Figure 4A**). Densitometric quantification of PD-L1 normalized to GAPDH and expressed relative to vehicle controls showed a significant increase in PD-L1 levels after *P. gingivalis* infection (mean +30% ± 14%; *p = .0361, two-tailed t-test; n = 3 independent experiments) (**Figure 4B**). Collectively, these results demonstrate that *P. gingivalis* reprograms glioma cells to adopt an immunosuppressive phenotype characterized by enhanced IL-6 secretion and PD-L1 expression. This dual activation mirrors pathogenic features observed in chronic *P. gingivalis* infection and periodontal disease, where sustained inflammation and PD-L1 induction enable local immune escape. In the context of GBM, such reprogramming may contribute to the establishment of an immunologically cold tumor microenvironment that blunts anti-tumor immune surveillance and facilitates tumor progression^24^. These observations are consistent with previous studies linking *P. gingivalis* infection to PD-L1 upregulation in epithelial and immune cell types^15,17,25,26^ and suggest that *P. gingivalis* may serve as a microbial trigger of the IL-6/PD-L1 axis in GBM pathogenesis.

**Figure 4:**
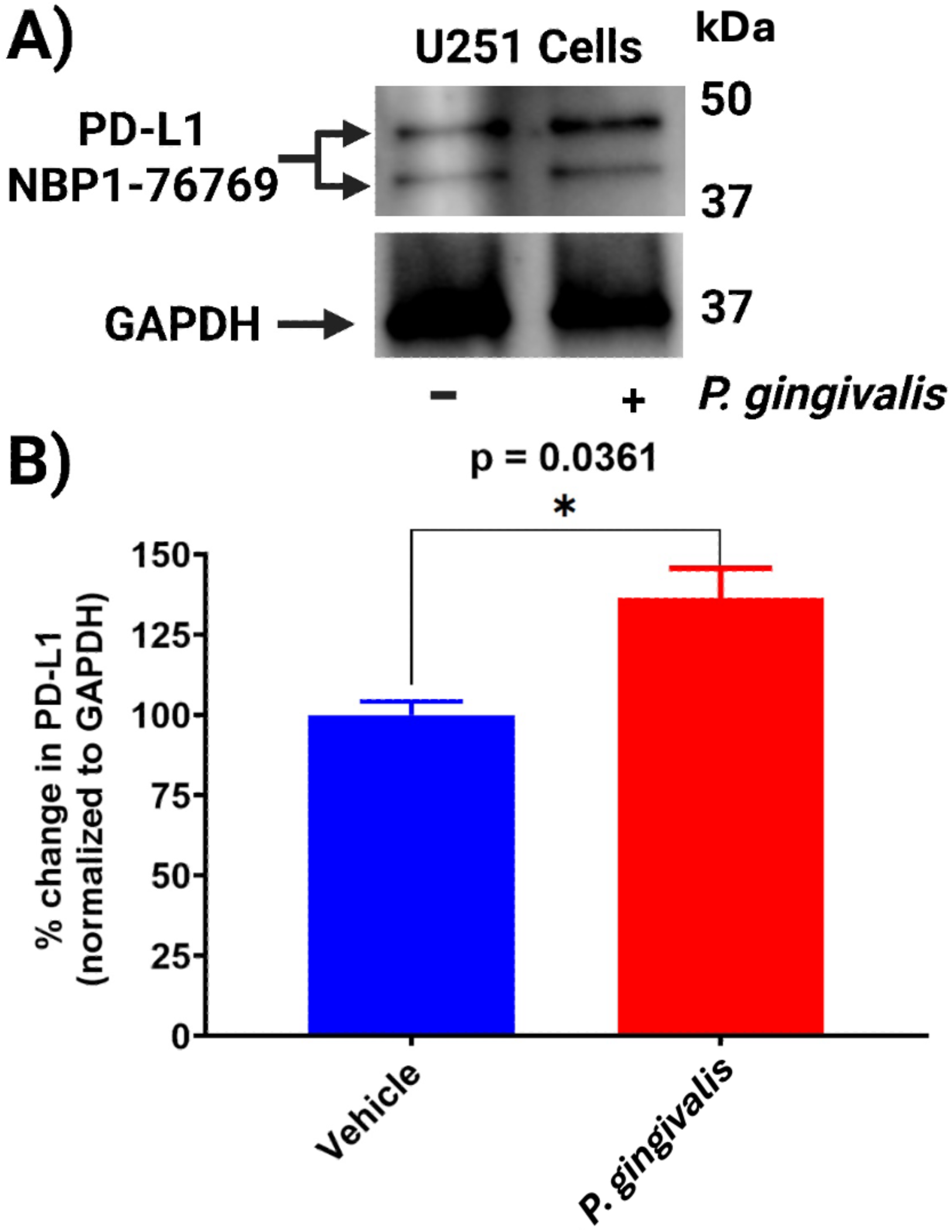
*P. gingivalis* increases PD-L1 expression in U251 glioma cells. **(A)** Immunoblot of U251 whole-cell lysates after exposure to vehicle (–) or *P. gingivalis* (+) for 24 h. Blots were probed with anti–PD-L1 (NBP1-76769); multiple bands correspond to PD-L1 glycoforms and isoforms. GAPDH was used as a loading control. Molecular mass markers (kDa) are indicated. **(B)** Densitometric quantification of PD-L1 normalized to GAPDH and expressed as % of vehicle control. *P. gingivalis* significantly increased PD-L1 levels (mean +30% ± 14%; *p = .0361, two-tailed t-test; *n* = 3 independent experiments). Bars show mean ± SD.

### P. gingivalis enhances the invasive capacity of U251 glioma cells

Having established that *P. gingivalis* stimulates IL-6 secretion and upregulates PD-L1 expression in glioma cells, which are hallmarks of a tumor-promoting and immunosuppressive microenvironment, we next aimed to determine whether *P. gingivalis* infection also affects the invasive behavior of glioma cells. Chronic IL-6 signaling and PD-L1 induction are both known to promote tumor cell motility, epi-thelial–mesenchymal transition (EMT), and extracellular matrix remodeling^27,28^; therefore, we hypothesize that *P. gingivalis*-mediated activation of these pathways could plausibly increase glioma cell invasiveness. To test this possibility, a Matrigel invasion assay was performed using U251 cells co-cultured with *P. gingivalis* (MOI 40). U251 glioma cells were seeded in the upper chamber of Matrigel-coated Transwell inserts containing serum-free medium supplemented with 0.5% FBS, while the lower chamber contained medium with 10% FBS as a chemoattractant. After 24 h of incubation, non-invading cells were removed, and cells that migrated through the Matrigel to the lower membrane surface were fixed and stained with crystal violet. A schematic overview of the experimental design is shown in **Figure 5A**. Representative images of crystal violet–stained membranes revealed minimal invasion in wells containing *P. gingivalis* alone, moderate baseline invasion by U251 cells, and markedly increased invasion when U251 cells were co-cultured with *P. gingivalis* (**Figure 5B**). A calibration curve demonstrated a strong linear correlation between absorbance at 590 nm and U251 cell number (R² = 0.9918), confirming the reliability of the crystal violet assay for quantification (**Figure 5C**). Our quantitative analysis showed a significant increase in the number of invasive U251 cells after exposure to *P. gingivalis* at an MOI of 40 compared with uninfected controls (**Figure 5D**).

**Figure 5:**
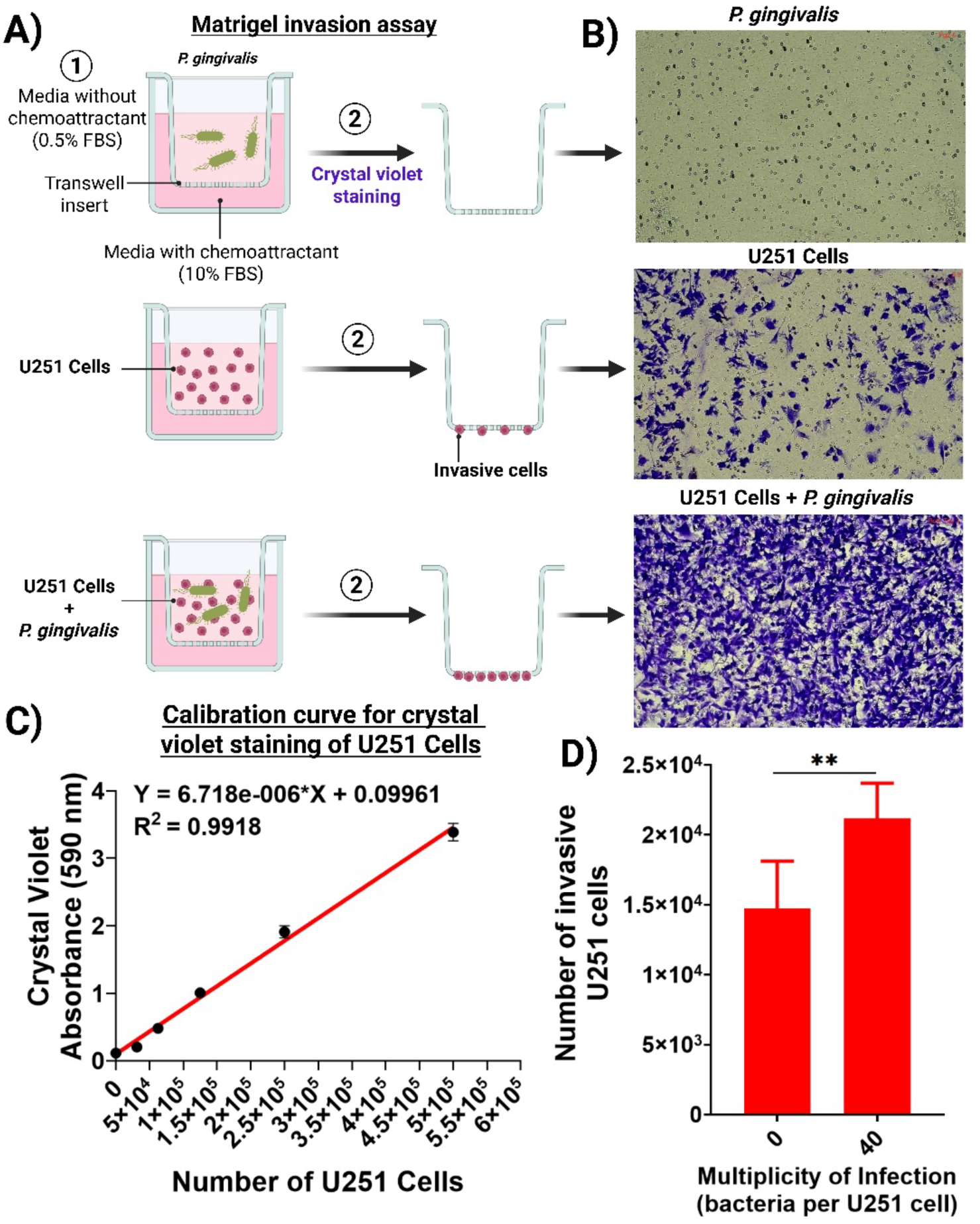
*P. gingivalis* increases the invasive ability of U251 glioma cells. **(A)** Schematic of the Matrigel invasion assay. U251 glioma cells were seeded in the top chamber of a Matrigel-coated transwell insert with or without *P. gingivalis* (MOI 40). The top chamber contained serum-free media with 0.5% FBS, while the bottom chamber had media with 10% FBS serving as a chemo-attractant. After 24 hours, non-invading cells were removed, and those that migrated through the Matrigel were stained with crystal violet. **(B)** Representative images of crystal violet-stained transwell membranes show minimal invasion with *P. gingivalis* alone (top), baseline invasion by U251 cells (middle), and increased invasion when U251 cells are co-cultured with *P. gingivalis* (bottom). **(C)** Calibration curve demonstrating the linear relationship (R² = 0.9918) between ab-sorbance at 590 nm and U251 cell number for invasion quantification using crystal violet staining. **(D)** Quantification indicates a significant increase in invasive U251 cells after exposure to *P. gin-givalis* at MOI 40 compared to uninfected control cells. Data are shown as mean ± SD; **p < .01 by unpaired two-tailed t-test.

To determine whether bacterial viability influences this pro-invasive phenotype, U251 cells were co-incubated with either live or heat-killed *P. gingivalis* at an MOI of 40 for 24 h. Quantification of invaded cells showed significantly higher invasion with live *P. gingivalis* compared to heat-killed bacteria (**Figure S1**). This indicates that bacterial viability is necessary for the full pro-invasive effect of *P. gingivalis* on glioma cells, implying that active bacterial processes, such as gingipain secretion, metabolic activity, or direct host–pathogen interactions, are crucial for promoting glioma cell invasiveness. Together, these results demonstrate that *P. gingivalis* not only reprograms glioma cells to produce IL-6 and express PD-L1 but also increases their invasiveness depending on viability.

## Discussion

This study supports a complex pathogenic model in which *P. gingivalis* infection contributes to both immune evasion and tumor aggressiveness, linking chronic microbial activity to GBM progression and recurrence (**Figure 6**). The findings of this study offer the first evidence that *P. gingivalis* gingipain virulence factors, RgpB and Kgp, are significantly elevated in GBM tissue samples compared to controls, with Kgp showing notably higher levels than RgpB. We show that *P. gingivalis* infection induces glioma cell IL-6 production, a cytokine linked to cancer resistance in GBM^22^, and upregulates PD-L1, known to promote GBM immune evasion^29^. We also demonstrate *in vitro* that *P. gingivalis* infection of U251 glioma cells results in significantly increased invasiveness compared to uninfected cells. Though foundational, these results are consistent with emerging evidence that *P. gingivalis* has oncogenic potential in body organs and tissues that it may translocate to after initial and during chronic infection of the oral cavity^30^.

**Figure 6:**
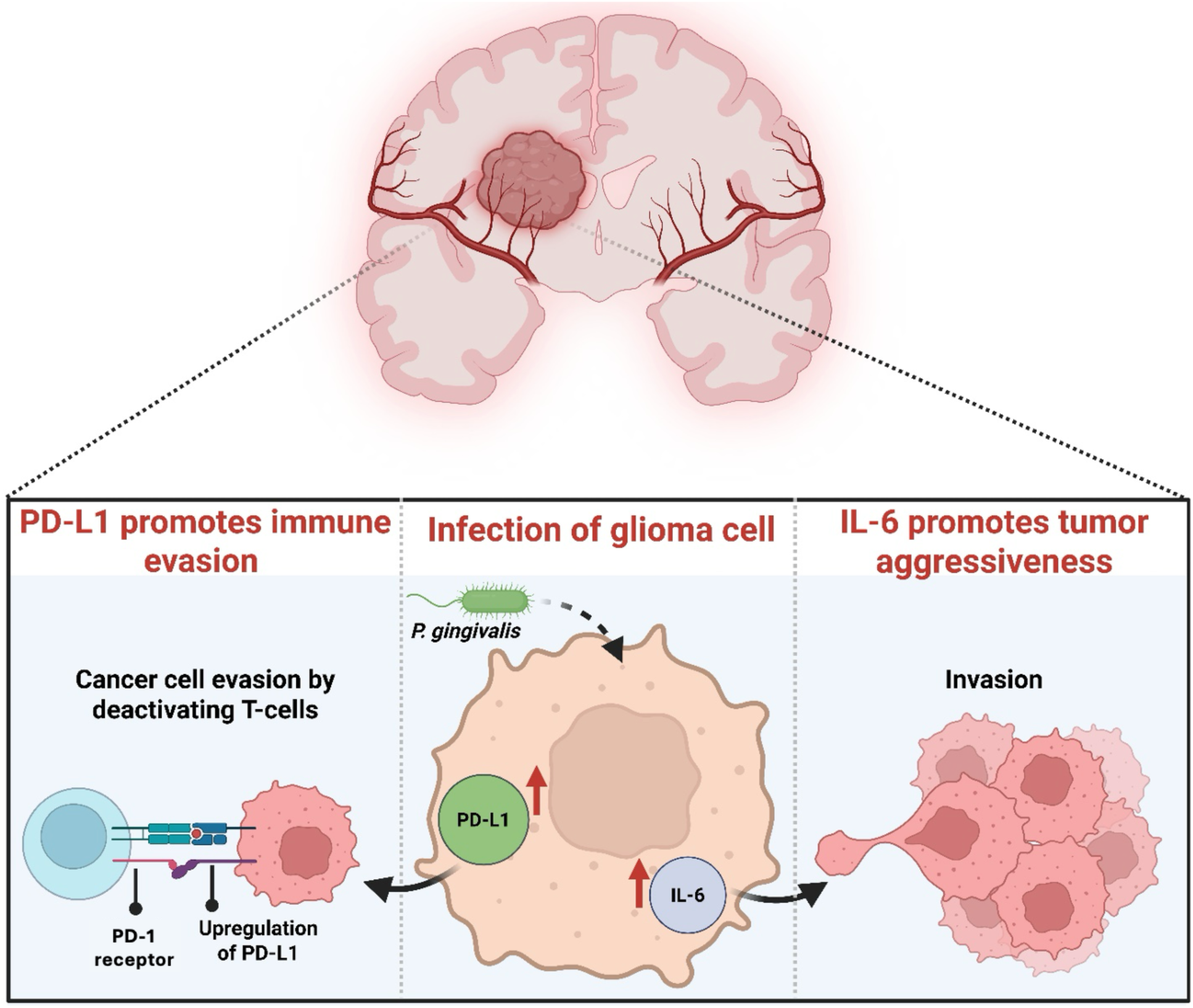
Proposed model of *P. gingivalis*–driven immune evasion and tumor aggressiveness in glioblastoma. Schematic representation of how *P. gingivalis* infection contributes to glioblastoma progression and recurrence through dual mechanisms. Upon infection of glioma cells, *P. gingivalis* induces upregulation of PD-L1 and IL-6. Elevated PD-L1 expression promotes immune evasion by suppressing cytotoxic T-cell activation, while increased IL-6 secretion enhances tumor aggressiveness by stimulating glioma cell invasion and proliferation. Together, these pathways establish a pro-tumorigenic and immunosuppressive microenvironment that facilitates glioblas-toma growth and recurrence.

Our results are also consistent with the hypothesis that GBM and AD may share a common underlying mechanism arising in the periphery^1^. As noted in the Introduction, members of our team have previously demonstrated elevated RgpB and Kgp antigens in AD brains^11^. An interesting difference between the current GBM study and the previous AD study is the relative distribution of the two gingipains. In AD brains, both RgpB and Kgp showed similarly high immunoreactivity levels. In contrast, GBM samples showed significantly higher Kgp staining compared to RgpB (****p < .0001) (**Figure 1D**), suggesting differential gingipain expression patterns between these two conditions.

Overexpression of Kgp compared to RgpB in the GBM tumor microenvironment could be a result of iron sequestration by GBM cells. Research has shown that GBM often overexpresses ferritin, sequestering iron within the tumor and creating a low iron microenvironment^31^. Kgp is the major iron scavenger for *P. gingivalis,* cleaving and liberating essential iron-containing heme from host proteins^32^, and its expression is increased in low iron environments resulting in increased pathogenic potential^33^. The difference between RgpB expression in GBM samples and AD brain samples could also be due to differences in *P. gingivalis* strains. Research has shown that *P. gingivalis* laboratory strains and clinical isolates from around the world exhibit different distributions of gingipains on the bacterial cell surface and secreted into the environment^34^. This raises the interesting possibility that certain strains of *P. gingivalis* may be oncogenic while other strains are more likely to drive neurodegeneration.

Another potential consequence of Kgp expression in the GBM tumor microenvironment is the disabling of antiviral defenses. Recent research has demonstrated that Kgp promotes viral infection by disabling the interferon pathway through proteolytic modification of major signaling components^35^. This mechanism may provide a basis for explaining the association of viruses, especially Herpes viruses such as HCMV, with GBM^6^.

Of note, two pediatric GBM samples from an 8-year-old and a 9-year-old showed detectable gingipain staining, with both samples showing Kgp immunoreactivity and one showing RgpB immunoreactivity **(Figure 2)**. GBM is uncommon in children, accounting for approximately 7-9% of central nervous system tumors, and has been observed to have similar histologic morphology between children and adults^36^. *P. gingivalis* is an age-related infection, increasing with age^37^, however, studies have shown that transmission between parents and young children can occur, even in infancy^38^.

Our results are supportive of a recent study that identified bacterial elements as part of the brain tumor microenvironment, with the finding of a greater number of intratumoral oral bacterial taxa compared to gut bacteria^39^. Along with several oral bacteria, the researchers identified intracellular *Porphyromonas* at the genus level in a set of tumor samples using Spatial Molecular Imaging (SIM)^39^. Our identification of gingipain antigens in GBM samples indicates that *Porphyromonas gingivalis* may be present at the species level, because gingipains are unique to *Porphyromonas gingivalis* in humans. The only other *Porphyromonas* species known to produce gingipains is *Porphyromonas gulae,* commonly found in companion animals such as dogs^40^.

This study has several limitations, the most significant being the relatively small sample size. A much larger sample size, with GBM samples from different age groups and geographic regions, is needed to determine if the expression of gingipains in GBM tumors is a common phenomenon. Additionally, the lack of genetic background information and incomplete treatment histories, such as prior radiation or chemotherapy exposure, limit our ability to fully interpret the observed associations. Future investigations should include lower-grade gliomas, such as grade I and II astrocytomas and grade III gliomas, to assess whether gingipain expression emerges early during gliomagenesis, or is restricted to high-grade disease.

While our current analyses focus on gingipain antigen detection, we are actively generating genomic and functional data to confirm *P. gingivalis* presence and viability within GBM tissue.

Ongoing work includes qPCR/ddPCR for *P. gingivalis*-specific genes, FISH-ISH co-localization, and culture-independent viability assays. To enhance physiological relevance, we are developing *ex vivo* GBM slice and organoid models that more accurately reflect tumor microenvironments and refining infection parameters through dose–response studies. Complementary mechanistic studies employing gingipain inhibitors, gingipain-deficient *P. gingivalis* mutants, and recombinant Kgp/RgpB will test whether IL-6 and PD-L1 upregulation are necessary and/or sufficient to drive invasion. Finally, murine *in vivo* studies will assess the impact of *P. gingivalis* infection on GBM progression, completing a translational framework that connects microbial detection to disease mechanism and therapeutic potential.

In conclusion, this study provides evidence that gingipains from *P. gingivalis* are expressed in GBM tumors and that *P. gingivalis* infection can increase the invasiveness of glioma cells. The reasons for this are currently unclear, as well as the mechanism for *P. gingivalis* and/or gingipain translocation from a peripheral *P. gingivalis* periodontal infection to the brain. One of the most intriguing findings of the current study was the observation that Kgp staining levels were markedly higher than RgpB in GBM samples, a pattern that differs from AD brains where both gingipains showed similarly high immunoreactivity^11^. More studies are needed to determine whether targeting *P. gingivalis* and its gingipains specifically can affect GBM progression.

A study of the role of LL-37 in preventing GBM progression may be an interesting future direction, as this natural human host defense peptide has been shown to inhibit the migration and clonogenicity of U251 glioma cells^41^. Importantly, *P. gingivalis*’s virulence factors can degrade the human host defense peptide LL-37^42^, possibly abrogating its anti-cancer and pro-autophagic activity. Moreover, one cannot exclude the possibility that modification and inactivation of LL-37 by citrullination can be executed by peptidyl arginine deiminase (PAD) expressed by *P. gingivalis*, the only PAD ortholog identified in prokaryotes^43^. Therefore, the degradation and/or modification of LL-37 by *P. gingivalis* is one possible contributing etiology of glioma. High-precision Kgp inhibitors are currently being tested in human clinical trials for the treatment of mild-to-moderate AD^44^, and if more preclinical research indicates *P. gingivalis* involvement in the pathogenesis of GBM, this same approach could be used in GBM clinical trials with biomarker evidence of Kgp expression in GBM biopsy samples.

**Figure S1:**
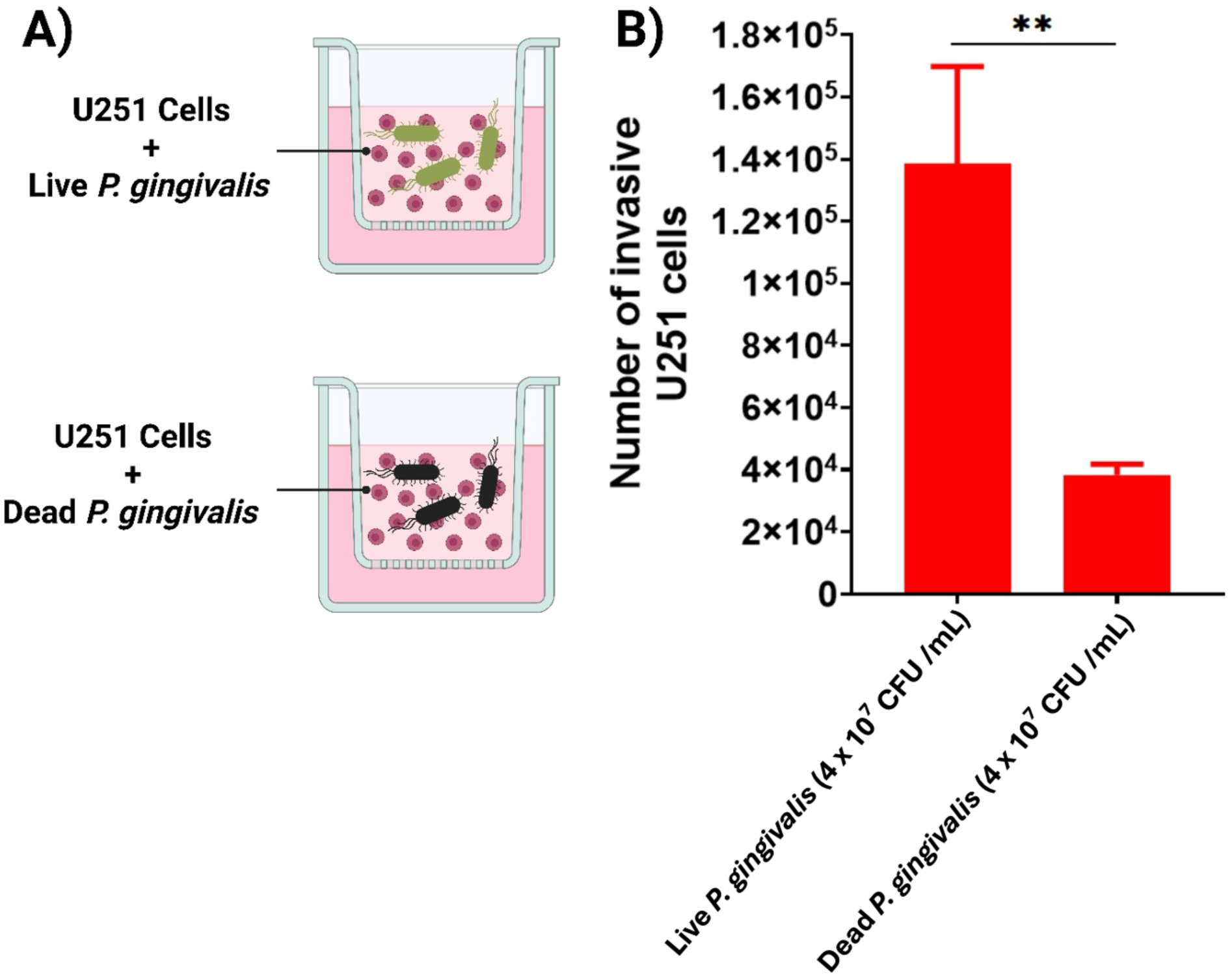
Bacterial viability increases the pro-invasive effect of *P. gingivalis* on U251 glioma cells. **(A)** Diagram of the Matrigel transwell invasion assay where U251 cells were co-incubated for 24 hours with live (top) or non-viable (heat-killed) *P. gingivalis* at 4×10^7^ CFU/mL (MOI 40)**. (B)** Quantification of invaded U251 cells shows significantly greater invasion with live *P. gingivalis* compared to dead *P. gingivalis* (mean ± SD; **p < .01, unpaired two-tailed t-test; n = 3 experiments). This suggests that bacterial viability is necessary for the maximum pro-invasive impact on glioma cells.

## Methods

### Immunohistochemistry

To investigate the frequency of gingipain antigens in glioblastoma tissue, immunohistochemistry (IHC) was performed on three 35-patient glioblastoma microarrays with 70 cores, two samples per patient (GL805-L51; Tissuearray.com, USA). One microarray was stained with hematoxylin and no primary antibody. The second microarray was stained with hematoxylin and CAB101, a primary antibody targeting arginine-gingipain B (RgpB)^11^. The third microarray was stained with hematoxylin and CAB102, a primary antibody targeting lysine-gingipain (Kgp)^11^. All other staining conditions were identical among the three arrays. An identical protocol was replicated on healthy tissue arrays (GLN801; Tissuearray.com, USA). 24 cores from 12 control patients with a pathology diagnosis of cerebrum gray matter or cerebrum white matter were included in the analysis. 10 cores in the glioblastoma microarray from the 5 control patients were included in the control tissue analysis, resulting in a sample size of 34 control cores from 17 patients.

### Immunohistochemical Image Analysis

Images were captured at 40X magnification and examined using ImageJ v1.54g. Color deconvolution (ColorDeconvolution2, H DAB setting, 8-bit transmittance) was performed to isolate DAB staining. A fixed threshold of 0–125 was consistently applied to all images, and the percentage of stained area was calculated. The background signal from the control array (no primary antibody stain) was subtracted from each corresponding sample.

### Cell Culture

Human glioma U251 cells (ECACC #89181493; Salisbury, UK) were maintained in DMEM supplemented with 10% FBS and 1% penicillin-streptomycin at 37 °C with 5% CO_2_. The cells were subcultured every 5 days, and passages 5–10 were used for experiments. Human astrocyte cells (ScienCell #1800; Carlsbad, CA, USA) were maintained in Astrocyte Medium (ThermoFisher #1261301; Waltham, MA, USA) at 37 °C with 5% CO_2_. The cells were subcultured every 5 days, and passages 5–10 were used for experiments.

### Cultivation of P. gingivalis Strain W83

*P. gingivalis* strain W83 was cultured under strictly anaerobic conditions using an AS150 Gloveless Anaerobic Chamber (Coy Laboratory Products, USA). The chamber atmosphere contained 80% nitrogen, 10% carbon dioxide, and 10% hydrogen, with oxygen levels maintained below 0.1% as verified by a resazurin indicator. Cultivation was performed in Brain Heart Infusion (BHI) Broth supplemented with hemin and vitamin K (Avantor Sciences, Cat. No. 89407-388), a medium formulated for the growth of fastidious anaerobic bacteria. A frozen stock of *P. gingivalis* W83 stored at –80 °C in 20% glycerol was thawed and streaked onto BHI agar plates containing 5% sheep blood (Hardy Diagnostics, Cat. No. A20). Plates were incubated anaerobically at 37 °C for 5 days, producing black-pigmented colonies characteristic of *P. gingivalis* due to heme accumulation. A single colony was subsequently inoculated into 5 mL of BHI broth with hemin and vitamin K and incubated statically at 37 °C for 48–72 hours, until the culture reached mid-log phase (OD₆₀₀ ≈ 0.8–1.0).

### P. gingivalis dosage determination

To determine a non-toxic inoculation dose of *P. gingivalis*, U251 cells were seeded in 48-well plates and incubated with varying CFU/mL for 24 h. Cell viability was assessed via MTT assay, and the highest non-lethal dose (4×10^7^ CFU/mL, MOI 40) was selected for subsequent experiments.

### IL-6 Assay

U251 cells and HA cells were seeded in 48-well plates and infected with *P. gingivalis* at varying concentrations under anaerobic conditions for 24 h. After infection, the supernatant was replaced with fresh medium, and cells were incubated for another 24 h. IL-6 levels in the supernatant were measured using ELISA (Thermo Fisher #EH21L6; Waltham, MA, USA) following the manufacturer’s instructions. Absorbance was read at 450 nm with a SpectraMax plate reader. Cell viability was confirmed by MTT assay.

### Matrigel Invasion Assay

Cell invasion was assessed using Corning BioCoat Matrigel Invasion Chambers (8.0 μm, #354480). U251 cells (1×10⁶ cells/chamber) were seeded in serum-free DMEM in the upper chamber, while lower chambers contained DMEM supplemented with 10% FBS. After 24 h, non-migrated cells were removed, and migrated cells were fixed and stained with 0.1% crystal violet. Three conditions were tested: U251 only, *P. gingivalis* + U251 (MOI 40), and *P. gingivalis* only. Chambers were incubated in DMSO (500 μL / well) for 30 min. Absorbance was measured at 590 nm, and the number of migrated cells was determined using the standard curve **(Figure 5C)** to convert absorbance values to cell counts.

### Calibration Curve for Crystal Violet Staining of U251 cells

U251 cells were plated in 48-well plates at serial two-fold dilutions starting from 5×10⁵ cells/well. After 4 h attachment, cells were fixed with 4% PFA, stained with 0.1% crystal violet (12 min), washed, and solubilized in 500 μL DMSO for 30 min. Absorbance was read at 590 nm, and cell number was plotted against absorbance to generate a standard curve.

### PD-L1 Assay

U251 cells were infected with *P. gingivalis* (4×10⁷ CFU/mL, MOI 40) under anaerobic conditions for 24 h, then trypsinized and analyzed for PD-L1 via Western blotting as described by Groeger et al^17^. Band intensities were quantified in ImageJ v1.54g and the PD-L1:GAPDH ratio was calculated for each sample and averaged per group (**Figure 4B**).

### Statistics

All experiments were conducted with independent biological replicates that were repeated, and statistical analyses were based on these biological replicates. Biological replicates are noted in each figure panel or legend. While no statistical methods were used to determine the sample sizes beforehand, the sizes in this study are comparable to or larger than those in prior reports. Data distribution was assumed normal, but this was not formally confirmed. Investigators were blinded during the analysis of experiments. The tissue samples used in this study were treated equally. Cells were randomly assigned to experimental conditions. No clinical, molecular, or cellular data points were excluded from the analyses. Bars indicate means, and error bars show the standard deviation (SD). Results were compared using t-tests in GraphPad. Generally, statistical significance is indicated with asterisks (*p ≤ .05, **p ≤ .01, ***p ≤ .001, ****p ≤ .0001), with exact p-values provided in figure panels or legends whenever possible.

## Data availability

Stained tissue images that support the findings of this study will be made available upon request.

## Code availability

No custom software, tools, or packages were used. The open-source software, tools, and packages used for data analysis in this study are referenced in the methods where applicable and include ColorDeconvolution (v2).

## Acknowledgements

The authors thank the Gunnar Heinrich Foundation for funding this work. They also acknowledge funding from an NIH Director’s Pioneer Award, Grant # 1DP1 OD029517, the SENS Foundation (now Lifespan Research Institute), Stanford University’s Discovery Innovation Fund, the Cisco University Research Program Fund, and Dr. James J. Truchard and the Truchard Foundation. We acknowledge Ms. Dagny C. Reese for her helpful discussions regarding the invasion assay protocol, and Dr. Jennifer S. Lin for her assistance in editing the manuscript.

## Author contributions

All authors made substantial contributions to the conception or design of the study; the acquisition, analysis, or interpretation of data; or drafting or revising the manuscript. All authors approved the manuscript. All authors agree to be personally accountable for individual contributions and to ensure that questions related to the accuracy or integrity or any part of the work are appropriately investigated and resolved and the resolution documented in the literature. EMM, LAB, SSD, and AEB conceived and designed the study. EMM and JC performed immunohistochemical experiments with guidance from FE. ZMMT cultured and maintained the *P. gingivalis* stock. EMM and JC cultured the U251 cells. EMM performed the *P. gingivalis* dosage determination. EMM performed the Matrigel invasion assays. LAB, JC, and EMM conducted the IL-6 studies. For the PD-L1 experiments, EMM created the experimental conditions, and JC ran the Western Blot. LAB conducted the statistical analysis with EMM. LAB developed the figures. The study was supervised by AEB, LAB, SSD, FE, and MIR. The manuscript was prepared by EMM, SSD, and LAB with input from all authors.

